# FasTag: automatic text classification of unstructured medical narratives

**DOI:** 10.1101/429720

**Authors:** Arturo Lopez Pineda, Oliver J. Bear, Guhan R. Venkataraman, Ashley M. Zehnder, Sandeep Ayyar, Rodney L. Page, Carlos D. Bustamante, Manuel A. Rivas

## Abstract

**Objective:** Unstructured clinical narratives are continuously being recorded as part of delivery of care in electronic health records, and dedicated tagging staff spend considerable effort manually assigning clinical codes for billing purposes; despite these efforts, label availability and accuracy are both suboptimal.

**Materials and Methods:** In this retrospective study, we trained long short-term memory (LSTM) recurrent neural networks (RNNs) on 52,722 human and 89,591 veterinary records. We investigated the accuracy of both separate-domain and combined-domain models and probed model portability. We established relevant baselines by training Decision Trees (DT) and Random Forests (RF), and using MetaMap Lite, a clinical natural language processing tool.

**Results:** We show that the LSTM-RNNs accurately classify veterinary and human text narratives into top-level categories with an average weighted macro F_1_ score of 0.74 and 0.68 respectively. In the “neoplasia” category, the model built with veterinary data has a high accuracy in veterinary data, and moderate accuracy in human data, with F_1_ scores of 0.91 and 0.70 respectively. Our LSTM method scored slightly higher than that of the DT and RF models.

**Discussion:** The use of LSTM-RNN models represents a scalable structure that could prove useful in cohort identification for comparative oncology studies.

**Conclusion:** Digitization of human and veterinary health information will continue to be a reality, particularly in the form of unstructured narratives. Our approach is a step forward for these two domains to learn from, and inform, one another.

## 1. OBJECTIVE

The increasing worldwide adoption of electronic health records (EHRs) has created numerous clinical narratives that are now stored in clinical databases. However, given the nature of the medical enterprise, a big portion of the data being recorded is in the form of unstructured clinical notes. Cohort studies, a form of cross-sectional studies that sample a group of patients with similar clinical characteristics, require quality phenotype labels, oftentimes not readily available alongside these notes.

In place of such labeling, diagnostic codes are the most common surrogates to true phenotypes. In clinical practice, dedicated tagging staff assign clinical codes to diagnoses either from the International Classification of Diseases (ICD) [1] or the Systematized Nomenclature of Medicine (SNOMED) after reading patient summaries. However, this time-consuming, error-prone task leads to only 60–80% of the assigned codes reflecting actual patient diagnoses [2], misjudgment of severity of conditions, and/or omission of codes altogether. For example, the relative inaccuracy of oncological medical coding [3–6] affects the quality of cancer registries [7] and cancer prevalence calculations [8–10]. Poorly-defined cancer types and poorly-trained coding staff who overuse the “not otherwise specified” code when classifying text exacerbate the problem.

Challenges in clinical coding also exist in veterinary medicine in the United States, where neither clinicians nor medical coders regularly apply diagnosis codes to veterinary visits. There are few incentives for veterinary clinicians to annotate their records; a lack of 1) a substantial veterinary third-party payer system and 2) legislation enforcing higher standards of veterinary EHRs (the U.S. Health Information Technology for Economic and Clinical Health Act of 2009 sets standards for human EHRs) compound the problem. Billing codes are thus rarely applicable across veterinary institutions unless hospitals share the same management structure and records system; even then, hospital-specific modifications exist. Less than five academic veterinary centers of a total of thirty veterinary schools in the United States have dedicated medical coding staff to annotate records using SNOMED-CT-Vet [11], a veterinary extension of SNOMED-CT constructed by the American Animal Hospital Association (AAHA) and maintained by the Veterinary Terminology Services Laboratory at the Virginia-Maryland Regional College of Veterinary Medicine [12].

The vast majority of veterinary clinical data is stored as free-text fields with very low rates of formal data curation, making data characterization a tall order. Further increasing variance in the data, veterinary patients come from many different environments, including hospitals [13], practices [14], zoos [15], wildlife reserves [16], army facilities [17], research facilities [18], breeders, dealers, exhibitors [19], livestock farms, and ranches [20]. It is thus important that a general method, agnostic of patient environment, is able to categorize veterinary EHRs for cohort identification solely based on free-text.

Automatic text classification is an emerging field that uses a combination of tools such as human medical coding, rule-based systems queries [21], natural language processing (NLP), statistical analyses, data mining, and machine learning (ML) [22]. In a previous study [23], we have shown the feasibility of automatic annotation of veterinary clinical narratives across a broad range of diagnoses with minimal preprocessing, but further exploration is needed to probe what we can learn from human-veterinary comparisons. Automatically adding meaningful disease-related tags to human and veterinary clinical notes using the same machinery would be a huge step forward in that exploration and could facilitate cross-species findings downstream.

Said integration has the potential to improve both veterinary and human coding accuracy as well as comparative analyses across species. Comparative oncology, for example, has accelerated the development of novel human anti-cancer therapies through the study of companion animals [24], especially dogs [25–28]. The National Institute of Health recently funded a multi-center institution called the Arizona Cancer Evolution Center (ACE) that aims to integrate data from a broad array of species to understand the evolutionarily conserved basis for oncology. As this group utilizes animal clinical and pathology data to identify helpful traits like species-specific cancer resistance, they would greatly benefit from improved cohort discovery through automated record tagging.

Veterinary schools across the United States (15 out of 30) have formed partnerships with their respective medical schools in order to perform cross-species translational research within the Clinical and Translational Science Award One Health Alliance (COHA, [29]). Of these schools, only two have active programs to assign disease codes to their medical records. The data for the rest represents the very use case of automatic text classification.

## 2. BACKGROUND AND SIGNIFICANCE

Automatic medical text classification aims to reduce the human burden of handling unstructured narratives. These computational natural language processing (NLP) methods can be divided into two groups: a) semantic processing and subsequent ML; and b) deep learning.

### Semantic processing and subsequent ML

These methods range from simple dictionary-based keyword-matching techniques and/or direct database queries to tools capable of interpreting the semantics of human language through lemmatization (removal of inflectional word endings), part-of-speech tagging, parsing, sentence breaking, word segmentation, and entity recognition [30]. Building the underlying dictionaries and manually crafting the rules that capture these diverse lexical elements both require time and domain expertise.

There is a growing interest in medical concept classification for clinical text; as such, many domain-specific semantic NLP tools (with various objectives, frameworks, licensing conditions, source code availabilities, language supports, and learning curves) have been developed for the medical setting. Such tools include MedLEE [31], MPLUS [32], MetaMap [33], KMCI [34], SPIN [35], HITEX [36], MCVS [37], ONYX [38], MedEx [39], cTAKES [40], pyConTextNLP [41], Topaz [42], TextHunter [43], NOBLE [44], and CLAMP [45]. However, there is no single NLP tool that can handle the broad problem of general medical classification. Instead, each method solves specific problems and applies its unique set of constraints.

ML downstream of the methods above requires featurization (concept extraction into columns and subsequent feature selection) in order to characterize text narratives in a machine-readable way. This can be done via term frequency-inverse document frequency (tf-idf), other vectorization techniques like Word2Vec [46], or manually curated rules. Semantic processing and downstream ML models have been shown to achieve high classification accuracy in human [47,48] and veterinary [49] free-text narratives for diseases well represented in training datasets (e.g. diabetes, influenza, and diarrhea). Additional success has been achieved in overall classification of clinical narratives with Decision Trees (DTs), Random Forests (RFs), and Support Vector Machines (SVMs) [50].

### Deep learning

Deep learning (DL) methods eliminate the need of feature engineering, harmonization, or rule creation. They learn hierarchical feature representations from raw data in an end-to-end fashion, requiring significantly less domain expertise than traditional machine-learning approaches [51].

DL is quickly emerging in the literature as a viable alternative method to traditional ML for the classification of clinical narratives, even in situations where limited labeled data is available [50]. The technique can help in the recognition of a limited number of categories from biomedical text [52,53]; identify psychiatric conditions of patients based on short clinical histories [54]; and accurately classify whether or not radiology reports indicate pulmonary embolism [55,56] whilst outperforming baseline methods (e.g. RFs or DTs). Previous studies have shown the possibility of using DL to label clinical narratives with medical subspecialties [57] (e.g. cardiology or neurology) or medical conditions [58] (e.g. advanced cancer or chronic pain), outperforming concept-extraction based methods. Furthermore, the use of DL to analyze clinical narratives has also facilitated the prediction of relevant patient attributes, such as in-hospital mortality, 30-day unplanned readmission, prolonged length of stay, and final discharge diagnosis [59].

### Significance:Text classification of human and veterinary medical records

Traditional NLP methods boast interpretability and flexibility but come at the steep cost of data quality control, formatting, normalization, domain knowledge, and time needed to generate meaningful heuristics (which oftentimes are not even generalizable to other datasets). Automatic text classification using deep learning is thus a logical choice to bypass these steps, classifying medical narratives from EHRs by leveraging supervised deep learning on big data. We expect that our efforts could facilitate rapid triaging of documents and cohort identification for biosurveillance.

## 3. METHODS

### Study Design

This retrospective cross-sectional chart review study uses medical records collected routinely as part of clinical care from two clinical settings: the veterinary teaching hospital at Colorado State University (CSU) and the Medical Information Mart for Intensive Care (MIMIC-III) from the Beth Israel Deaconess Medical Center in Boston, Massachusetts [60]. Both datasets were divided in two smaller datasets - training datasets containing 70% of the original datasets (used to build TensorFlow [61] deep learning models), and validation datasets containing 30% of the original datasets. We measured the accuracy of the models, calculating the F_1_ score of each top-level disease category.

For comparison, we investigated the effect of using MetaMap [33], a NLP tool that extracts clinically-relevant terms, on the accuracy of our models. We also explored the possibility of out-of-domain generalization, testing the MIMIC-trained model on the CSU validation data and vice versa (and ran separate tests for MetaMapped versions, as well). Finally, we investigated the effect of merging the MIMIC and CSU training datasets to test the efficacy of data augmentation. Figure 1 shows a diagram of our study design. Our code to run all models can be found in a public repository (https://github.com/rivas-lab/FasTag).

**Figure 1.**
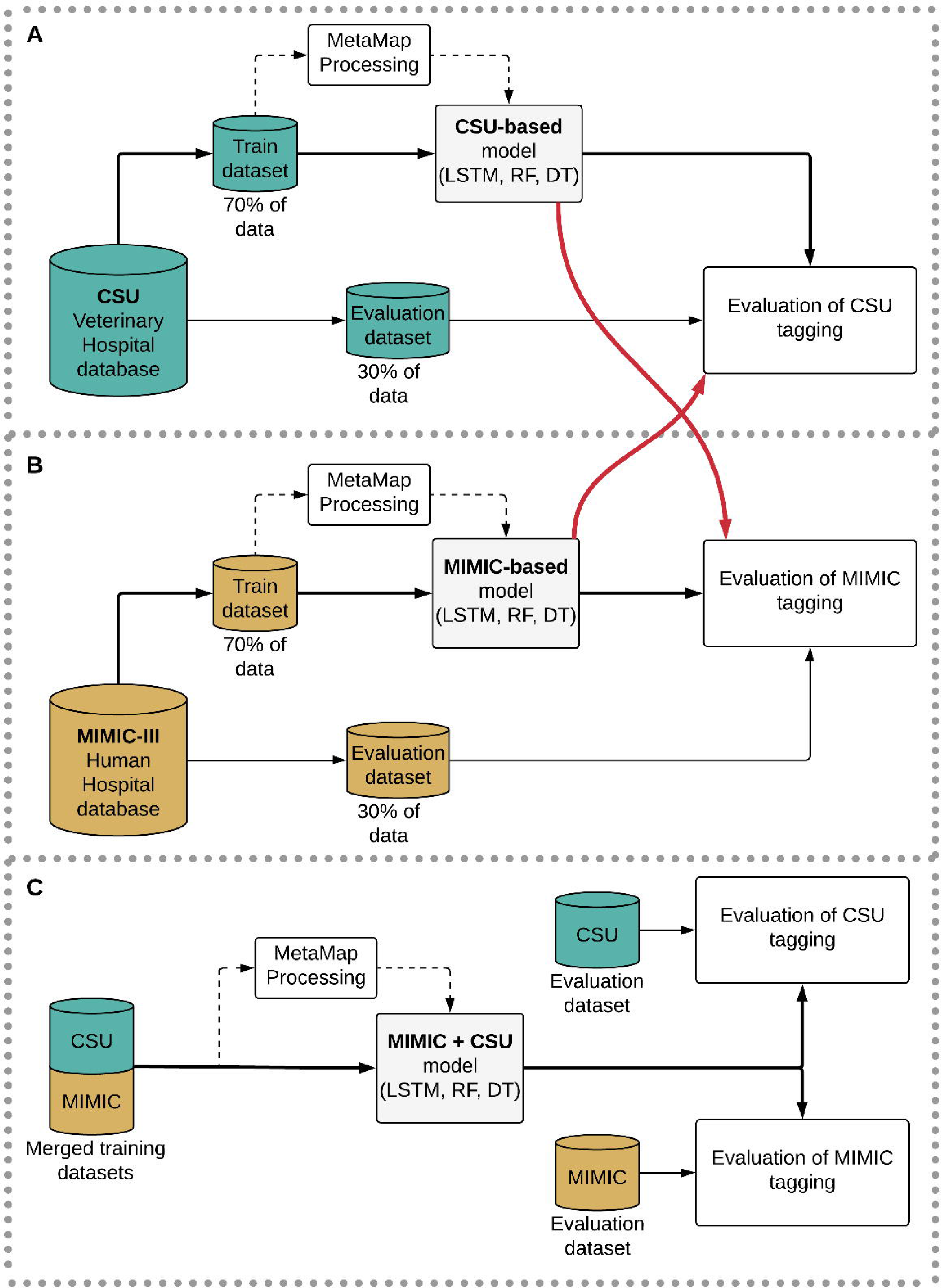
Diagram of the training and evaluation design. Relevant acronyms: MIMIC: Medical Information Mart for Intensive Care; CSU: Colorado State University; MetaMap, a tool for recognizing medical concepts in text; LSTM: long-short term memory recurrent neural network classifier; RF: Random Forest classifier; DT: Decision Tree classifier

### Clinical Settings

#### Veterinary Medical Hospital at Colorado State University (CSU)

This is a tertiary care referral teaching hospital with inpatient and outpatient facilities, serving all specialties of veterinary medicine. After consultation, veterinarians enter patient information into a custom-built veterinary EHR, including structured fields such as entry and discharge dates, patient signalment (species, breed, age, sex, etc.), and SNOMED-CT-Vet codes. There are also options to input free-text clinical narratives with various sections including history, assessment, diagnosis, prognosis, and medications. These records are subsequently coded; the final diagnostic codes represent single or multiple specific diagnoses or post-coordinated expressions (a combination of two or more concepts).

#### Medical Information Mart for Intensive Care (MIMIC-III)

The Beth Israel Deaconess Medical Center is a tertiary care teaching hospital at Harvard Medical School in Boston, Massachusetts. The MIMIC-III database, a publicly available dataset which we utilize in this study, contains information on patients admitted to the critical care unit at the hospital [60]. We were interested in the free-text hospital discharge summaries in this database. These records are coded for billing purposes and have complete diagnoses per patient (the database is publicly available, and thus represents the best possible medical coding annotation scenario for a hospital). Free-text fields in this database contain no protected health information.

### Top level disease categories

Mapping between ICD and SNOMED codes is a challenging task but can promote semantic interoperability between our two domains. We used ICD top-level groups of diseases as the labels for the records that we aimed to extract. Table 1 shows the mapping between codes in ICD (versions 9 and 10) and SNOMED-CT (including the Veterinary extension), which was manually curated by two board-certified veterinarians trained in clinical coding (co-authors AZM, and RLP).

**Table 1.**
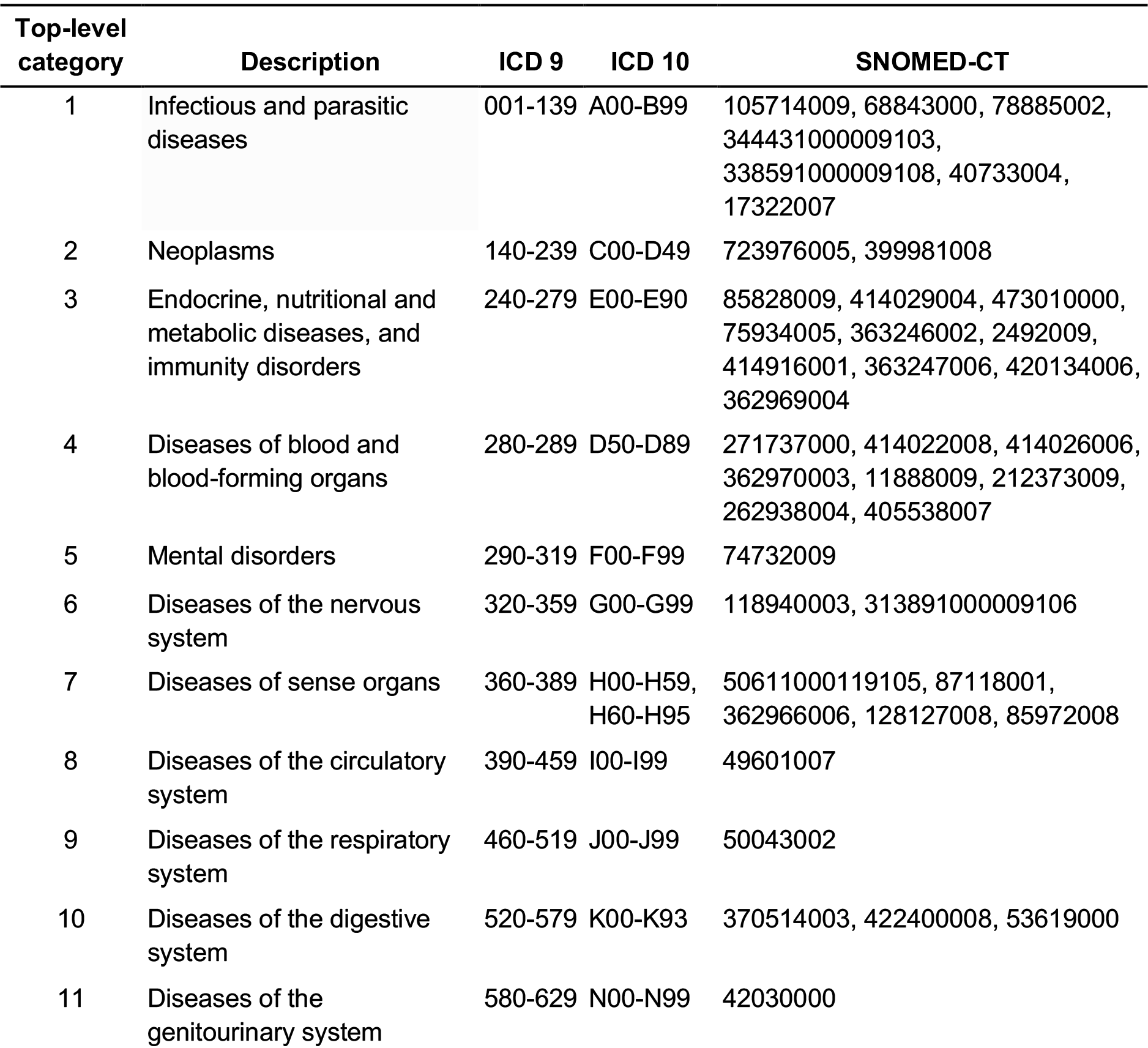
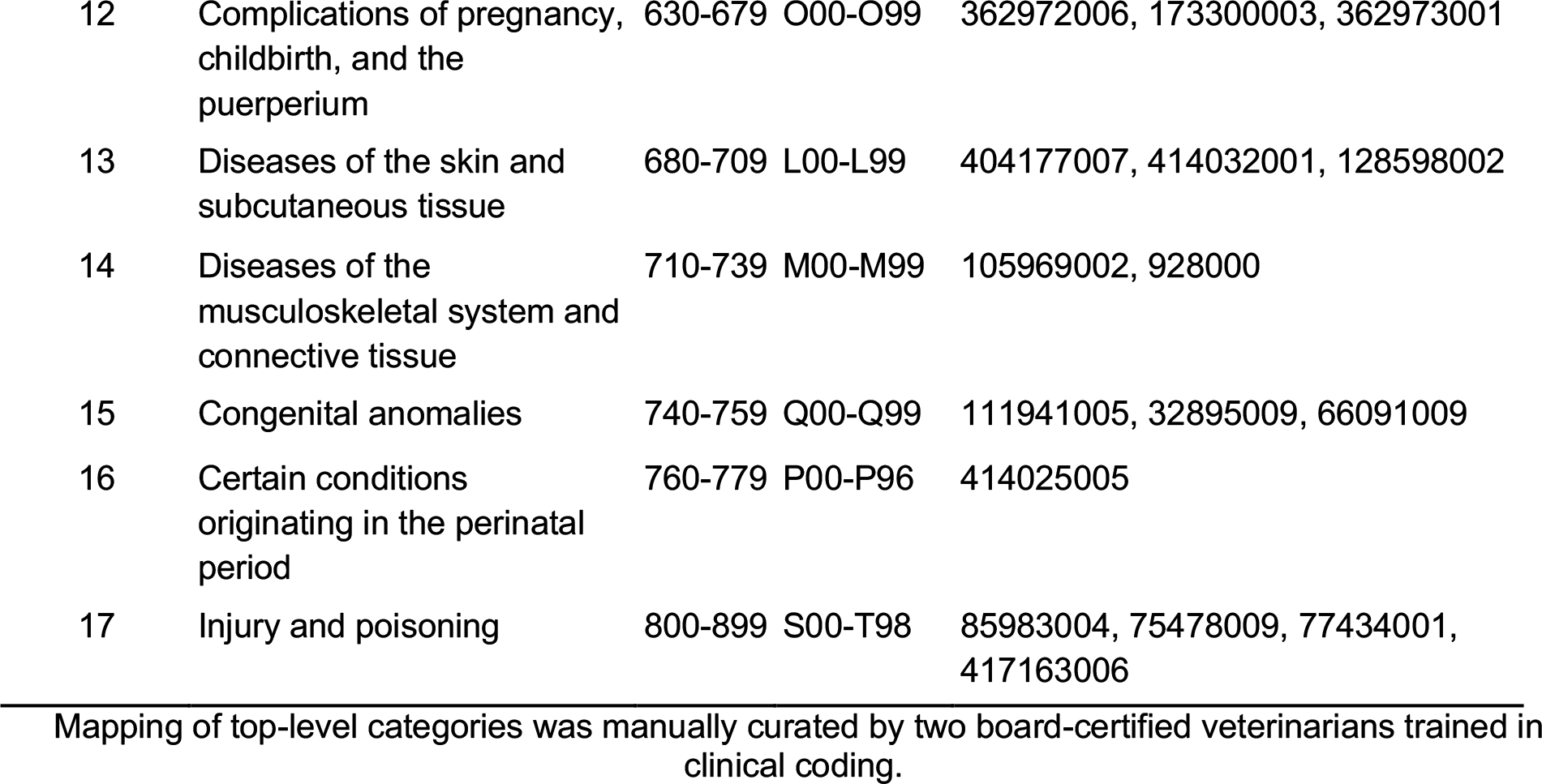
Top-level coding mapping between ICD 9, 10, and SNOMED-CT

### Deep learning

We chose a long short-term memory (LSTM) recurrent neural network (RNN) architecture (which is able to handle variable-length sequences while using previous inputs to inform current time steps) for this multi-label text classification task [62]. The LSTM shares parameters across time steps as it unrolls, which allows it to handle sequences of variable length. In this case, these sequences are a series of word “embeddings” (created by mapping specific words to corresponding numeric vectors) from clinical narratives. Words are represented densely (rather than sparsely, as in Bag-of-Words or tf-idf models) using the Global Vectors for Word Representation (GloVe) [63] word embeddings. These embeddings learn a vector space representation of words such that words with similar contexts appear in a similar vector space, and also capture global statistical features of the training corpus.

LSTMs have proven to be flexible enough to be used in many different tasks, such as machine translation, image captioning, medication prescription, and forecasting disease diagnosis using structured data [62]. The RNN can efficiently capture sequential information and theoretically model long-range dependencies, but empirical evidence has shown this is difficult to do in practice [64]. The sequential nature of text lends itself well to LSTMs, which have memory cells that can maintain information for over multiple time steps (words) and consist of a set of gates that control when information enters and exits memory, making them an ideal candidate architecture.

We first trained the model over a variety of hyperparameters for the model trained on MIMIC data and calculated the model’s validation accuracy over all combinations of them, finding the set of [learning rate = 0.001, dropout rate = 0.5, batch size = 256, training epochs = 100, hidden layer size = 200, LSTM layers =1] to be the optimal setting. We proceeded to use this hyperparameter set for all of the models trained, assuming that this set would be amenable to the task at hand regardless of training dataset. We then proceeded to train a set of six models on the MIMIC, CSU, and MIMIC+CSU data; one each in which MetaMap was used to map terms back to UMLS terms, and one each in which MetaMap was not. We finally determined F_1_ scores on the corresponding validation sets for each of these models.

### Baseline classifier comparisons

A combination of several NLP and ML models have similarly aimed to classify clinical narratives [50,65]. We selected two of these classifiers: DTs and RFs. DTs are ML models constructed around a branching boolean logic [66]. Each node in the tree can take a decision that leads to other nodes in a tree structure; there are no cycles allowed. The RF classifier is an ensemble of multiple decision trees created by randomly selecting samples of the training data. The final prediction is done via a consensus voting mechanism of the trees in the forest.

We featurized the narratives using tf-idf, a statistic that reflects word importance in the context of other documents in a corpus and a standard ML modeling strategy for representing text, to convert the narratives into a tabular format [50]. The hyperparameters of both baseline models, like the LSTM, were tuned on the validation set.

We used MetaMap Lite [67], a NLP tool which leverages the Unified Medical Language System (UMLS) Metathesaurus to identify SNOMED [68] or ICD [69] codes from clinical narratives. MetaMap’s algorithm includes five steps: 1) parsing of text into simple noun phrases; 2) variant generation of phrases to include all derivations of words (i.e. synonyms, acronyms, meaningful spelling variants, combinations, etc.); 3) candidate retrieval of all UMLS strings that contains at least one variant from the previous step; 4) evaluation and ranking of each candidate, mapping between matched term and the Metathesaurus concept using metrics of centrality, variation, coverage, and cohesiveness; 5) construction of complete mappings to include those mappings that are involved in disjointed parts of the phrase (e.g. ‘ocular’ and ‘complication’ can together be mapped to a single term, ‘ocular complication’). MetaMap incorporates the use of ConText [70], an algorithm for the identification of negation in clinical narratives. For additional information on how we used and evaluated MetaMap, please refer to Supplementary Material 1.

### Statistical analysis

#### Evaluation metric

For all models we trained (LSTM, DT, and RF), we used the same evaluation metrics previously reported for MetaMap Lite [67]: a) precision, defined as the proportion of documents which were assigned the correct category; b) recall, defined as the proportion of documents from a given category that were correctly identified; and c) F_1_ score, defined as the harmonic average of precision and recall. Formulas for these metrics are provided below:

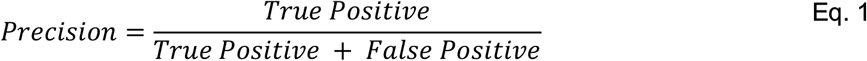

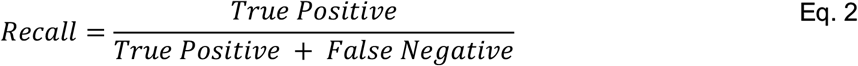

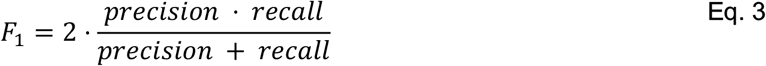

Our task is framed as a multi-label classification problem, where each approach predicts multiple top-level categories for each observation using a single model. In order to combine all class-specific F_1_ scores, we averaged the F_1_ score for each label, weighting the labels by their supports (the number of true instances for each label, to account for label imbalance).

#### Domain adaptation

The portability of trained algorithms on independent domains has previously been used as a metric of model robustness in systems that leverage NLP and machine learning [71]. We evaluated the ability of our trained LSTM models to be used in a cross-species context. We utilized the MIMIC-trained model to classify the medical records in the CSU database, and vice versa, assessing performance as before. We also assess the classifier trained on the combined training sets.

## 4. RESULTS

We investigated the application of deep learning to free-text unstructured clinical narratives on two cohorts: veterinary medical records from CSU, and human medical records in the MIMIC-III database. We show the evaluation of the deep learning models built using human and veterinary records, as well as the portability between them.

### Patients

The CSU dataset contains medical records from 33,124 patients and 89,591 hospital visits between February 2007 and July 2017. Patients encompassed seven mammalian species, including dogs (*Canis Lupus*, 80.8%), cats (*Felis Silvestris*, 11.4%), horses (*Equus Caballus*, 6.5%), cattle (*Bos Taurus*, 0.7%), pigs (*Sus Scrofa,* 0.3%), goats (*Capra hircus*, 0.2%), sheep (*Ovis Aries*, 0.1%), and other unspecified mammals (0.1%). In contrast, the MIMIC-III database contains medical records from 38,597 distinct human adult patients (aged 16 years or above) and 7,870 neonates admitted between 2001 and 2008, encompassing 52,722 unique hospital admissions to the critical care unit between 2001 and 2012. Table 2 summarizes the category breakdowns of both databases. Only those patients with a diagnosis in their record were considered.

**Table 2.** Database statistics of patients, records, and species (records with diagnosis).

|  | CSU | MIMIC |
| --- | --- | --- |
| **Medical Records** | N = 89,591 | N = 52,722 |
| Patients | 33,124 | 41,126 |
| Hospital Visits | 89,591 | 49,785 |
| **Species** |  |  |
| Humans (*Homo Sapiens*) | n.a. | 52,722 |
| Dogs (*Canis Lupus*) | 72,420 | n.a. |
| Cats (*Felis Silvestris*) | 10,205 | n.a. |
| Horses (*Equus Caballus*) | 5,819 | n.a. |
| Other mammals | 1,147 | n.a. |
| **Category** |  |  |
| 1. Infectious | 11,454 | 10,074 |
| 2. Neoplasia | 36,108 | 6,223 |
| 3. Endo-Immune | 17,295 | 24,762 |
| 4. Blood | 10,171 | 13,481 |
| 5. Mental | 511 | 10,989 |
| 6. Nervous | 7,488 | 9,168 |
| 7. Sense organs | 15,085 | 2,688 |
| 8. Circulatory | 8,733 | 30,054 |
| 9. Respiratory | 11,322 | 17,667 |
| 10. Digestive | 22,776 | 14,646 |
| 11. Genitourinary | 8,892 | 14,932 |
| 12. Pregnancy | 136 | 133 |
| 13. Skin | 21,147 | 4,241 |
| 14. Musculoskeletal | 22,921 | 6,739 |
| 15. Congenital | 3,347 | 2,334 |
| 16. Perinatal | 54 | 3,661 |
| 17. Injury | 9,873 | 16,121 |

### Evaluation of Deep Learning models

We trained deep-learning models (as well as DT/RF baselines) for the human, veterinary, and merged (human and veterinary) datasets and tested each on their own domain, as well as the other domains. We built our LSTM model using the Python programming language (version 2.7), TensorFlow [61] (version 1.9), and the ‘scikit-learn’ library (version 0.19.2) [72]. The training was performed on an Amazon® Deep Learning AMI, a cloud-based platform running the Ubuntu operating system with pre-installed CUDA dependencies. Average weighted macro F_1_ scores for models across all categories are shown in Table 3; a full list of F_1_ scores by category can be found in Supplementary Material 1. The “neoplasia” category results, which we found interesting, are shown in Table 4.

**Table 3.** Average F_1_ scores using various training and validation dataset combinations for all categories.

| Configuration |  |  | Model evaluation<br>(Weighted F<sub>1</sub> score) |  |  |
| --- | --- | --- | --- | --- | --- |
| Training | Validation | MetaMap | DT | RF | LSTM |
| MIMIC | MIMIC | No | 0.60 | 0.64 | <u>**0.65**</u> |
|  |  | Yes | 0.60 | 0.63 | <u>**0.70**</u> |
| CSU | CSU | No | 0.55 | 0.61 | <u>**0.72**</u> |
|  |  | Yes | 0.54 | 0.60 | <u>**0.75**</u> |
| MIMIC | CSU | No | 0.22 | 0.24 | <u>**0.28**</u> |
|  |  | Yes | 0.23 | 0.20 | <u>**0.31**</u> |
| CSU | MIMIC | No | <u>**0.31**</u> | 0.20 | 0.23 |
|  |  | Yes | 0.28 | 0.19 | <u>**0.36**</u> |
| MIMIC+CSU | CSU | No | 0.57 | 0.62 | <u>**0.67**</u> |
|  |  | Yes | 0.57 | 0.62 | <u>**0.76**</u> |
| MIMIC+CSU | MIMIC | No | 0.60 | <u>**0.63**</u> | 0.58 |
|  |  | Yes | 0.60 | <u>**0.63**</u> | 0.60 |
| MIMIC+CSU | MIMIC+CSU | No | 0.59 | 0.64 | <u>**0.68**</u> |
|  |  | Yes | 0.59 | 0.63 | <u>**0.71**</u> |
| Average |  |  | 0.489 | 0.506 | <u>**0.571**</u> |

Evaluation metrics for Decision Tree (DT), Random Forest (RF), and Long Short Term Memory Recurrent Neural Network (LSTM-RNN) on validation datasets with and without MetaMap term extraction. Bolded and underlined numbers represent the best scores for the specific configuration of training data, validation data, and MetaMap toggle.

**Table 4.** F_1_ scores using various training and validation dataset combinations for the “neoplasia” category.

| Configuration |  |  | Model evaluation<br>(Weighted F<sub>1</sub> score) |  |  |
| --- | --- | --- | --- | --- | --- |
| Training | Validation | MetaMap | DT | RF | LSTM |
| MIMIC | MIMIC | No | 0.39 | 0.45 | <u>**0.66**</u> |
|  |  | Yes | 0.4 | 0.45 | <u>**0.76**</u> |
| CSU | CSU | No | 0.81 | 0.86 | <u>**0.91**</u> |
|  |  | Yes | 0.8 | 0.86 | <u>**0.91**</u> |
| MIMIC | CSU | No | 0.3 | 0.53 | <u>**0.69**</u> |
|  |  | Yes | 0.45 | 0.37 | <u>**0.75**</u> |
| CSU | MIMIC | No | 0.46 | 0.58 | <u>**0.70**</u> |
|  |  | Yes | 0.5 | <u>**0.58**</u> | 0.54 |
| MIMIC+CSU | CSU | No | 0.74 | 0.8 | <u>**0.87**</u> |
|  |  | Yes | 0.74 | 0.8 | <u>**0.87**</u> |
| MIMIC+CSU | MIMIC | No | 0.4 | 0.47 | <u>**0.67**</u> |
|  |  | Yes | 0.42 | 0.45 | <u>**0.72**</u> |
| MIMIC+CSU | MIMIC+CSU | No | 0.81 | <u>**0.86**</u> | 0.85 |
|  |  | Yes | 0.81 | 0.86 | <u>**0.90**</u> |
| Average |  |  |  |  |  |

Evaluation metrics for the “neoplasia” category Decision Tree (DT), Random Forest (RF), and Long Short Term Memory Recurrent Neural Network (LSTM-RNN) on validation datasets with and without MetaMap term extraction. Bolded and underlined numbers represent the best scores for the specific configuration of training data, validation data, and MetaMap toggle.

## 5. DISCUSSION

Applying deep learning to unstructured free-text clinical narratives in electronic health records offers a relatively simple, low-effort means to bypass the traditional bottlenecks in medical coding. Circumventing the need for data harmonization was very important for the datasets, which contained a plethora of domain-and setting-specific misspellings, abbreviations, and jargon (these issues would have greatly impacted the performance of the LSTM and the NLP’s entity recognition). MetaMap was useful in this regard given its ability to parse clinical data, but much work is still needed to improve recognition of terms in veterinary and human domains.

There is moderate evidence of domain adaptation in the “neoplasia” category, with F_1_ scores of 0.69-0.70 (Table 4). This process involved training a model on the data in one database and testing on the data in the other, without fine-tuning. It is evident that the high classification accuracy (F_1_ score = 0.91) obtained by the CSU model in the neoplasia category is decreased when testing the same model on the MIMIC data. One possible explanation is the difference in clinical settings; CSU is a tertiary care veterinary hospital specializing in oncological care, and the clinical narratives that arise in a critical care unit like the MIMIC dataset do not necessarily compare. Moreover, the records were not coded in the same way, the clinicians did not receive the same training, and the documents apply to different species altogether. Despite these differences, however, our LSTM model was general enough to be able to accurately classify medical narratives at the top level of depth independently in both datasets. The achieved cross-domain accuracy is nonetheless encouraging. Given enough training data and similar-enough clinical narratives, one could conceivably imagine a general model that is highly effective across domains.

Models performed usually better on their respective validation datasets in those categories with more training samples. For example, the CSU-trained model (25,276 samples) had significantly better performance in the “neoplasia” category than the MIMIC-trained model (4,356 samples), while the MIMIC-trained model (21,038 samples) had better performance in the diseases of the circulatory system category than the CSU-trained model (6,133 samples).

The usefulness of even top-level characterizations in the veterinary setting cannot be understated; usually, a veterinarian must read the full, unstructured text in order to get any information about the patient they are treating. Rapid selection of documents with specific types of clinical narratives (such as oncological cases, which our model performed well on) could lead to better cohort studies for comparative research. The repeated use of a series of such LSTM models for subsequent, increasingly-specific classifications thus represents a scalable, hierarchical tagging structure that could prove extremely useful in stratifying patients by specific diseases, severities, and protocols.

## 6. CONCLUSION

In this era of increasing deployment of EHRs, it is important to provide tools that facilitate cohort identification. Our deep learning approach (LSTM model) was able to automatically classify medical narratives with minimal human preprocessing. In a future with enough training data, it is possible to foresee a scenario in which these models can accurately tag every clinical concept, regardless of data input. The expansion of veterinary data availability and the subsequently enormous potential of domain adaptation like we saw in the neoplasia category could prove to be exciting chapters in reducing bottlenecks in public health research at large; it is thus of critical importance to continue studying novel sources of data that can rapidly be used to augment classification models.

A reliable addition to existing rule-based and natural language processing strategies, deep learning is a promising tool for accelerating public health research.

## DECLARATIONS

### Data availability

Veterinary data presented here belongs to the Colorado State University, which may grant access to this data on a case-by-case basis to researchers who obtain the necessary Data Use Agreement (DUA) and IRB approvals.

Human data presented here belongs to the Beth Israel Deaconess Medical Center in Boston, Massachusetts, which can be accessed after signing a data usage agreement in the MIT Lab for Computational Physiology at https://mimic.physionet.org/

#### Acknowledgments

The authors wish to acknowledge Dr. Katie M. Kanagawa for her valuable support in editing this manuscript. Also, Devin Johnson, DVM, MS, for her contribution to clinical coding and comparison with coding from the MetaMap tool.

### Contributions

ALP, OBDW, GRV, and AMZ designed the study. RLP provided access to the veterinary data. OJBDW, GRV, ALP, AMZ, and SA extracted, formatted, and performed analysis of the data. AMZ, RLP, CDB and MAR provided interpretation of the results. ALP drafted the manuscript, and all authors contributed critically, read, revised and approved the final version.

### Ethics approval

This research was reviewed and approved by Stanford’s Institutional Review Board (IRB), which provided a non-human subject determination under eProtocol 46979. Consent was not required.

