## Supplementary Material 1 for "FasTag: automatic text classification of unstructured medical narratives"

### I. Evaluation of MetaMap on veterinary records

MetaMap Lite has not previously been applied to veterinary data. As such, we endeavored to verify that the software works as expected when applied in this context. Two board-certified veterinarians trained in clinical coding independently evaluated MetaMap-extracted terms from 19 randomly-selected records. Disagreements were resolved via in-depth discussion and consensus. Evaluation of the NLP was done using a confusion matrix for each record. The ground truth in this evaluation was considered to be the human experts; terms selected by MetaMap but not by reviewers were considered false positives, terms selected by reviewers but not by MetaMap were considered false negatives, and terms selected by both the reviewers and MetaMap were considered true positives. We assumed there were no true negatives. This process resulted in a weighted-average precision of 0.62, recall of 0.82, and $F_1$ score of 0.71 for the CSU data, as compared to a previously reported weighted-average precision of 0.67, recall of 0.53, and $F_1$ score of 0.58 for human clinical narratives. The results of this evaluation can be seen in Table A.

**Table A.** NLP (MetaMap) Evaluation.

| Record No. | True positive | False positive | False negative | Total | Recall | Precision | $F_1$-score |
|---|---|---|---|---|---|---|---|
| 1 | 29 | 3 | 11 | 43 | 0.73 | 0.91 | 0.81 |
| 2 | 86 | 23 | 66 | 175 | 0.57 | 0.79 | 0.66 |
| 3 | 103 | 11 | 103 | 217 | 0.50 | 0.90 | 0.64 |
| 4 | 208 | 43 | 81 | 332 | 0.72 | 0.83 | 0.77 |
| 5 | 96 | 18 | 38 | 152 | 0.72 | 0.84 | 0.77 |
| 6 | 55 | 19 | 20 | 94 | 0.73 | 0.74 | 0.74 |
| 7 | 127 | 22 | 124 | 273 | 0.51 | 0.85 | 0.64 |

| | | | | | | | |
|---|---|---|---|---|---|---|---|
| 8 | 50 | 16 | 34 | 100 | 0.60 | 0.76 | 0.67 |
| 9 | 97 | 25 | 79 | 201 | 0.55 | 0.80 | 0.65 |
| 10 | 66 | 5 | 26 | 97 | 0.72 | 0.93 | 0.81 |
| 11 | 84 | 28 | 58 | 170 | 0.59 | 0.75 | 0.66 |
| 12 | 32 | 8 | 26 | 66 | 0.55 | 0.80 | 0.65 |
| 13 | 112 | 29 | 93 | 234 | 0.55 | 0.79 | 0.65 |
| 14 | 156 | 27 | 88 | 271 | 0.64 | 0.85 | 0.73 |
| 15 | 61 | 6 | 14 | 81 | 0.81 | 0.91 | 0.86 |
| 16 | 124 | 24 | 53 | 201 | 0.70 | 0.84 | 0.76 |
| 17 | 114 | 37 | 51 | 202 | 0.69 | 0.75 | 0.72 |
| 18 | 108 | 24 | 47 | 179 | 0.70 | 0.82 | 0.75 |
| 19 | 126 | 30 | 100 | 256 | 0.56 | 0.81 | 0.66 |
| | | | | **Weighted averages** | **0.62** | **0.82** | **0.71** |

Comparison of reviewer and MetaMap term extractions for 19 records. True Positive (TP) = MetaMap correctly matched term to code; False Negative (FN) = MetaMap did not extract a term the experts did; False Positive (FP) = MetaMap matched a term not identified by the expert; Total = Total number of terms matched in that document.

The entities recognized by MetaMap included all semantic types described in the website documentation (https://metamap.nlm.nih.gov/Docs/SemanticTypes_2018AB.txt). We did not restrict the search to any particular semantic type. Evaluation of the NLP was done using a confusion matrix for each record. For the first example, the confusion matrix would be as shown in Table B.

**Table B.** Confusion Matrix for MetaMap Lite evaluation on veterinary data.

| | Human selected entities | Human fail to select entities | Total |
|---|---|---|---|
| MetaMap selected entities | 29 (true positive) | 3 (false positive) | 32 |
| MetaMap fail to select entities | 11 (false negative) | ?? (true negative) | 11 |
| Total | 40 | 3 | 43 |

In our current design, it is not possible to know how many entities were missed by both the MetaMap tagger and the two board-certified veterinarians with clinical coding training. We assumed this number to be zero. Given this assumption, the evaluation of MetaMap in veterinary narratives is comparable to previously reported evaluations in human narratives.

### MetaMap results

After MetaMap selected terms and assigned concept-unique identifiers (CUI), we only retained the matched term. The order in which terms appear in the original text was also preserved. Finally, we constructed MetaMap-matched clinical narratives, where those terms assigned "False" by the ConText algorithm are added a "no_" prefix. All terms were converted to lowercase, special characters (not alphanumeric) were removed, and extraneous spaces were simplified to single underscores. When multiple matched CUIs existed, duplicates of the word were kept in the new clinical narrative. This process was conducted independently for each document before passed as input into the deep learning model. Table C shows an example of one free-text clinical narrative processed with MetaMap.

**Table C.** Example of free-text and MetaMap-extracted veterinary record

| Free-text clinical narrative | MetaMap-matched clinical terms |
|---|---|
| [PET_NAME], a 2 year old female spayed Bernese Mountain Dog, presented for evaluation of allergic dermatitis and otitis. | year old female bernese_mountain_dog evaluation allergic_dermatitis otitis. |
| [PET_NAME] had a history of recurrent UTI's as a puppy, but these problems have resolved. | history recurrent_utis puppy problems. |
| [PET_NAME] has a 1 year history of recurrent otitis externa and pruritus directed primarily at her feet and ears. | year history recurrent_otitis otitis_externa pruritus feet ears. |
| The pruritus and otitis problems appear to be more severe during the winter. | pruritus otitis problems more severe winter. |

| | |
|---|---|
| She also has a two month history of recurrent conjunctivitis that first developed after another dog in the house returned from a kennel. | month history recurrent conjunctivitis first dog house. |

A 2-year old female dog patient with recurrent otitis and allergic dermatitis. Both the narrative (left) and the MetaMapped version (right) show that the treatment included prednisone (among other important clinical details). For the purposes of this manuscript, the pet and owner's name were manually de-identified

### II. Classification performance across categories

**Table D.** Using Long Short Term Memory (LSTM) Recurrent Neural Network (RNN)

| | **MIMIC → MIMIC** | | **CSU → CSU** | | **MIMIC → CSU** | | **CSU → MIMIC** | | **MIMIC + CSU → MIMIC + CSU** | | **MIMIC + CSU → MIMIC** | | **MIMIC + CSU → CSU** | |
|---|---|---|---|---|---|---|---|---|---|---|---|---|---|---|
| **Category Name** | **Full** | **MM** | **Full** | **MM** | **Full** | **MM** | **Full** | **MM** | **Full** | **MM** | **Full** | **MM** | **Full** | **MM** |
| Infectious and parasitic diseases | 0.61 | 0.69 | 0.54 | 0.61 | 0.26 | 0.28 | 0.48 | 0.34 | 0.58 | 0.66 | 0.56 | 0.64 | 0.53 | 0.63 |
| Neoplasms | 0.66 | 0.76 | 0.91 | 0.91 | 0.69 | 0.75 | 0.70 | 0.54 | 0.87 | 0.87 | 0.67 | 0.72 | 0.85 | 0.90 |
| Endocrine, nutritional and metabolic diseases, and immunity disorders | 0.86 | 0.86 | 0.69 | 0.71 | 0.33 | 0.29 | 0.15 | 0.31 | 0.74 | 0.77 | 0.69 | 0.70 | 0.61 | 0.71 |
| Diseases of blood and blood-forming organs | 0.56 | 0.56 | 0.81 | 0.81 | 0.23 | 0.32 | 0.30 | 0.08 | 0.55 | 0.62 | 0.37 | 0.39 | 0.62 | 0.82 |
| Mental disorders | 0.38 | 0.59 | 0.00 | 0.00 | 0.01 | 0.01 | 0.00 | 0.00 | 0.22 | 0.41 | 0.22 | 0.27 | 0.00 | 0.00 |
| Diseases of the nervous system | 0.50 | 0.56 | 0.65 | 0.70 | 0.24 | 0.25 | 0.04 | 0.41 | 0.59 | 0.64 | 0.58 | 0.53 | 0.62 | 0.71 |
| Diseases of sense organs | 0.00 | 0.00 | 0.81 | 0.82 | 0.00 | 0.00 | 0.28 | 0.15 | 0.71 | 0.75 | 0.15 | 0.40 | 0.77 | 0.82 |

| | | | | | | | | | | | | | | |
|---|---|---|---|---|---|---|---|---|---|---|---|---|---|---|
| Diseases of the circulatory system | 0.94 | 0.94 | 0.62 | 0.67 | 0.20 | 0.28 | 0.50 | 0.80 | 0.88 | 0.88 | 0.87 | 0.90 | 0.63 | 0.67 |
| Diseases of the respiratory system | 0.71 | 0.81 | 0.67 | 0.71 | 0.23 | 0.32 | 0.37 | 0.65 | 0.72 | 0.76 | 0.73 | 0.74 | 0.63 | 0.71 |
| Diseases of the digestive system | 0.66 | 0.77 | 0.73 | 0.77 | 0.40 | 0.43 | 0.00 | 0.35 | 0.74 | 0.77 | 0.67 | 0.68 | 0.71 | 0.77 |
| Diseases of the genitourinary system | 0.73 | 0.75 | 0.65 | 0.70 | 0.30 | 0.34 | 0.00 | 0.19 | 0.66 | 0.70 | 0.58 | 0.66 | 0.63 | 0.70 |
| Complications of pregnancy, childbirth, and the puerperium | 0.00 | 0.00 | 0.00 | 0.64 | 0.00 | 0.00 | 0.10 | 0.00 | 0.00 | 0.61 | 0.00 | 0.16 | 0.00 | 0.68 |
| Diseases of the skin and subcutaneous tissue | 0.09 | 0.20 | 0.74 | 0.75 | 0.00 | 0.01 | 0.03 | 0.07 | 0.66 | 0.69 | 0.29 | 0.28 | 0.59 | 0.76 |
| Diseases of the musculoskeletal system and connective tissue | 0.05 | 0.22 | 0.71 | 0.75 | 0.01 | 0.02 | 0.00 | 0.18 | 0.60 | 0.60 | 0.32 | 0.07 | 0.68 | 0.74 |
| Congenital anomalies | 0.05 | 0.04 | 0.09 | 0.25 | 0.01 | 0.08 | 0.31 | 0.01 | 0.09 | 0.29 | 0.01 | 0.07 | 0.04 | 0.37 |
| Certain conditions originating in the perinatal period | 0.97 | 0.95 | 0.00 | 0.00 | 0.00 | 0.01 | 0.38 | 0.00 | 0.79 | 0.79 | 0.44 | 0.94 | 0.00 | 0.00 |
| Injury and poisoning | 0.60 | 0.66 | 0.65 | 0.68 | 0.36 | 0.34 | 0.17 | 0.20 | 0.54 | 0.56 | 0.43 | 0.39 | 0.63 | 0.66 |
| **Macro F1** | 0.49 | 0.55 | 0.54 | 0.62 | 0.19 | 0.22 | 0.22 | 0.25 | 0.58 | 0.67 | 0.45 | 0.50 | 0.50 | 0.63 |
| **Weighted Macro F1** | 0.65 | 0.70 | 0.72 | 0.75 | 0.28 | 0.31 | 0.23 | 0.36 | 0.68 | 0.71 | 0.58 | 0.60 | 0.67 | 0.76 |

**Table E.** Using Decision Trees (DT)

| | MIMIC → MIMIC | | CSU → CSU | | MIMIC → CSU | | CSU → MIMIC | | MIMIC + CSU → MIMIC + CSU | | MIMIC + CSU → MIMIC | | MIMIC + CSU → CSU | |
|---|---|---|---|---|---|---|---|---|---|---|---|---|---|---|
| **Category Name** | **Full** | **MM** | **Full** | **MM** | **Full** | **MM** | **Full** | **MM** | **Full** | **MM** | **Full** | **MM** | **Full** | **MM** |
| Infectious and parasitic diseases | 0.51 | 0.51 | 0.41 | 0.40 | 0.16 | 0.21 | 0.28 | 0.27 | 0.46 | 0.45 | 0.50 | 0.51 | 0.41 | 0.40 |
| Neoplasms | 0.39 | 0.40 | 0.81 | 0.80 | 0.30 | 0.45 | 0.46 | 0.50 | 0.74 | 0.74 | 0.40 | 0.42 | 0.81 | 0.81 |
| Endocrine, nutritional and metabolic diseases, and immunity disorders | 0.78 | 0.77 | 0.47 | 0.46 | 0.32 | 0.30 | 0.43 | 0.32 | 0.67 | 0.68 | 0.77 | 0.78 | 0.49 | 0.49 |
| Diseases of blood and blood-forming organs | 0.59 | 0.60 | 0.63 | 0.62 | 0.21 | 0.14 | 0.17 | 0.15 | 0.61 | 0.60 | 0.58 | 0.58 | 0.63 | 0.63 |
| Mental disorders | 0.43 | 0.44 | 0.05 | 0.05 | 0.01 | 0.01 | 0.01 | 0.01 | 0.43 | 0.43 | 0.44 | 0.44 | 0.05 | 0.03 |
| Diseases of the nervous system | 0.42 | 0.41 | 0.39 | 0.36 | 0.16 | 0.15 | 0.26 | 0.24 | 0.40 | 0.41 | 0.39 | 0.40 | 0.42 | 0.44 |
| Diseases of sense organs | 0.09 | 0.11 | 0.67 | 0.66 | 0.02 | 0.03 | 0.13 | 0.14 | 0.58 | 0.57 | 0.09 | 0.09 | 0.67 | 0.68 |
| Diseases of the circulatory system | 0.89 | 0.88 | 0.35 | 0.35 | 0.19 | 0.18 | 0.38 | 0.34 | 0.80 | 0.80 | 0.89 | 0.89 | 0.41 | 0.41 |
| Diseases of the respiratory system | 0.66 | 0.67 | 0.43 | 0.40 | 0.23 | 0.22 | 0.43 | 0.42 | 0.58 | 0.59 | 0.66 | 0.66 | 0.47 | 0.45 |
| Diseases of the digestive system | 0.57 | 0.58 | 0.53 | 0.52 | 0.35 | 0.31 | 0.49 | 0.41 | 0.54 | 0.53 | 0.53 | 0.53 | 0.54 | 0.53 |
| Diseases of the genitourinary system | 0.64 | 0.65 | 0.43 | 0.43 | 0.21 | 0.19 | 0.27 | 0.27 | 0.59 | 0.59 | 0.66 | 0.66 | 0.45 | 0.44 |
| Complications | 0.04 | 0.06 | 0.09 | 0.12 | 0.00 | 0.00 | 0.00 | 0.00 | 0.04 | 0.13 | 0.07 | 0.00 | 0.07 | 0.16 |

| | | | | | | | | | | | | | | |
|---|---|---|---|---|---|---|---|---|---|---|---|---|---|---|
| of pregnancy, childbirth, and the puerperium | | | | | | | | | | | | | | |
| Diseases of the skin and subcutaneous tissue | 0.21 | 0.22 | 0.58 | 0.58 | 0.25 | 0.09 | 0.16 | 0.10 | 0.51 | 0.51 | 0.22 | 0.20 | 0.59 | 0.58 |
| Diseases of the musculoskeletal system and connective tissue | 0.27 | 0.25 | 0.56 | 0.55 | 0.13 | 0.21 | 0.23 | 0.22 | 0.48 | 0.47 | 0.25 | 0.26 | 0.56 | 0.57 |
| Congenital anomalies | 0.05 | 0.07 | 0.16 | 0.17 | 0.03 | 0.04 | 0.04 | 0.06 | 0.15 | 0.16 | 0.07 | 0.06 | 0.19 | 0.22 |
| Certain conditions originating in the perinatal period | 0.90 | 0.90 | 0.00 | 0.00 | 0.06 | 0.01 | 0.00 | 0.00 | 0.65 | 0.62 | 0.83 | 0.81 | 0.00 | 0.00 |
| Injury and poisoning | 0.54 | 0.54 | 0.41 | 0.38 | 0.22 | 0.20 | 0.26 | 0.20 | 0.52 | 0.52 | 0.54 | 0.54 | 0.46 | 0.46 |
| **Macro F1** | 0.47 | 0.47 | 0.41 | 0.40 | 0.17 | 0.16 | 0.24 | 0.21 | 0.51 | 0.52 | 0.46 | 0.46 | 0.42 | 0.43 |
| **Weighted Macro F1** | 0.60 | 0.60 | 0.55 | 0.54 | 0.22 | 0.23 | 0.31 | 0.28 | 0.59 | 0.59 | 0.60 | 0.60 | 0.57 | 0.57 |

**Table F.** Using Random Forests (RF)

| | **MIMIC → MIMIC** | | **CSU → CSU** | | **MIMIC → CSU** | | **CSU → MIMIC** | | **MIMIC + CSU → MIMIC + CSU** | | **MIMIC + CSU → MIMIC** | | **MIMIC + CSU → CSU** | |
|---|---|---|---|---|---|---|---|---|---|---|---|---|---|---|
| **Category Name** | **Full** | **MM** | **Full** | **MM** | **Full** | **MM** | **Full** | **MM** | **Full** | **MM** | **Full** | **MM** | **Full** | **MM** |
| Infectious and parasitic diseases | 0.55 | 0.55 | 0.44 | 0.43 | 0.07 | 0.09 | 0.22 | 0.10 | 0.50 | 0.50 | 0.54 | 0.55 | 0.43 | 0.44 |
| Neoplasms | 0.45 | 0.45 | 0.86 | 0.86 | 0.53 | 0.37 | 0.58 | 0.58 | 0.80 | 0.80 | 0.47 | 0.45 | 0.86 | 0.86 |
| Endocrine, nutritional and metabolic diseases, and | 0.83 | 0.83 | 0.50 | 0.50 | 0.34 | 0.33 | 0.19 | 0.13 | 0.74 | 0.74 | 0.83 | 0.83 | 0.55 | 0.53 |

| | | | | | | | | | | | | | | |
|---|---|---|---|---|---|---|---|---|---|---|---|---|---|---|
| immunity disorders | | | | | | | | | | | | | | |
| Diseases of blood and blood-forming organs | 0.66 | 0.65 | 0.71 | 0.71 | 0.23 | 0.21 | 0.06 | 0.06 | 0.67 | 0.67 | 0.65 | 0.65 | 0.72 | 0.72 |
| Mental disorders | 0.44 | 0.43 | 0.00 | 0.01 | 0.02 | 0.03 | 0.00 | 0.00 | 0.40 | 0.42 | 0.43 | 0.43 | 0.00 | 0.00 |
| Diseases of the nervous system | 0.39 | 0.38 | 0.45 | 0.45 | 0.16 | 0.13 | 0.21 | 0.23 | 0.41 | 0.41 | 0.37 | 0.37 | 0.47 | 0.51 |
| Diseases of sense organs | 0.02 | 0.02 | 0.76 | 0.75 | 0.00 | 0.00 | 0.08 | 0.09 | 0.67 | 0.66 | 0.03 | 0.02 | 0.75 | 0.76 |
| Diseases of the circulatory system | 0.93 | 0.93 | 0.36 | 0.36 | 0.19 | 0.18 | 0.17 | 0.20 | 0.86 | 0.86 | 0.93 | 0.93 | 0.44 | 0.44 |
| Diseases of the respiratory system | 0.72 | 0.72 | 0.48 | 0.45 | 0.27 | 0.24 | 0.36 | 0.42 | 0.66 | 0.65 | 0.71 | 0.71 | 0.53 | 0.52 |
| Diseases of the digestive system | 0.62 | 0.62 | 0.57 | 0.56 | 0.39 | 0.34 | 0.36 | 0.38 | 0.58 | 0.57 | 0.57 | 0.56 | 0.58 | 0.58 |
| Diseases of the genitourinary system | 0.70 | 0.71 | 0.50 | 0.49 | 0.26 | 0.27 | 0.18 | 0.20 | 0.64 | 0.65 | 0.71 | 0.71 | 0.48 | 0.51 |
| Complications of pregnancy, childbirth, and the puerperium | 0.00 | 0.08 | 0.08 | 0.12 | 0.00 | 0.00 | 0.00 | 0.00 | 0.03 | 0.03 | 0.00 | 0.00 | 0.04 | 0.08 |
| Diseases of the skin and subcutaneous tissue | 0.16 | 0.16 | 0.65 | 0.64 | 0.00 | 0.00 | 0.11 | 0.09 | 0.58 | 0.57 | 0.17 | 0.15 | 0.66 | 0.65 |
| Diseases of the musculoskeletal system and connective tissue | 0.18 | 0.16 | 0.62 | 0.61 | 0.03 | 0.03 | 0.15 | 0.19 | 0.51 | 0.50 | 0.16 | 0.17 | 0.63 | 0.63 |
| Congenital anomalies | 0.04 | 0.04 | 0.12 | 0.09 | 0.00 | 0.00 | 0.00 | 0.01 | 0.15 | 0.15 | 0.04 | 0.03 | 0.20 | 0.19 |
| Certain conditions | 0.97 | 0.95 | 0.00 | 0.00 | 0.00 | 0.00 | 0.00 | 0.00 | 0.79 | 0.78 | 0.93 | 0.93 | 0.00 | 0.00 |

| | | | | | | | | | | | | | | |
|---|---|---|---|---|---|---|---|---|---|---|---|---|---|---|
| originating in the perinatal period | | | | | | | | | | | | | | |
| Injury and poisoning | 0.58 | 0.56 | 0.48 | 0.40 | 0.30 | 0.22 | 0.16 | 0.06 | 0.57 | 0.56 | 0.56 | 0.55 | 0.55 | 0.53 |
| **Macro F1** | 0.48 | 0.48 | 0.45 | 0.44 | 0.16 | 0.14 | 0.17 | 0.16 | 0.56 | 0.56 | 0.48 | 0.47 | 0.46 | 0.47 |
| **Weighted Macro F1** | 0.64 | 0.63 | 0.61 | 0.60 | 0.24 | 0.20 | 0.20 | 0.19 | 0.64 | 0.63 | 0.63 | 0.63 | 0.62 | 0.62 |

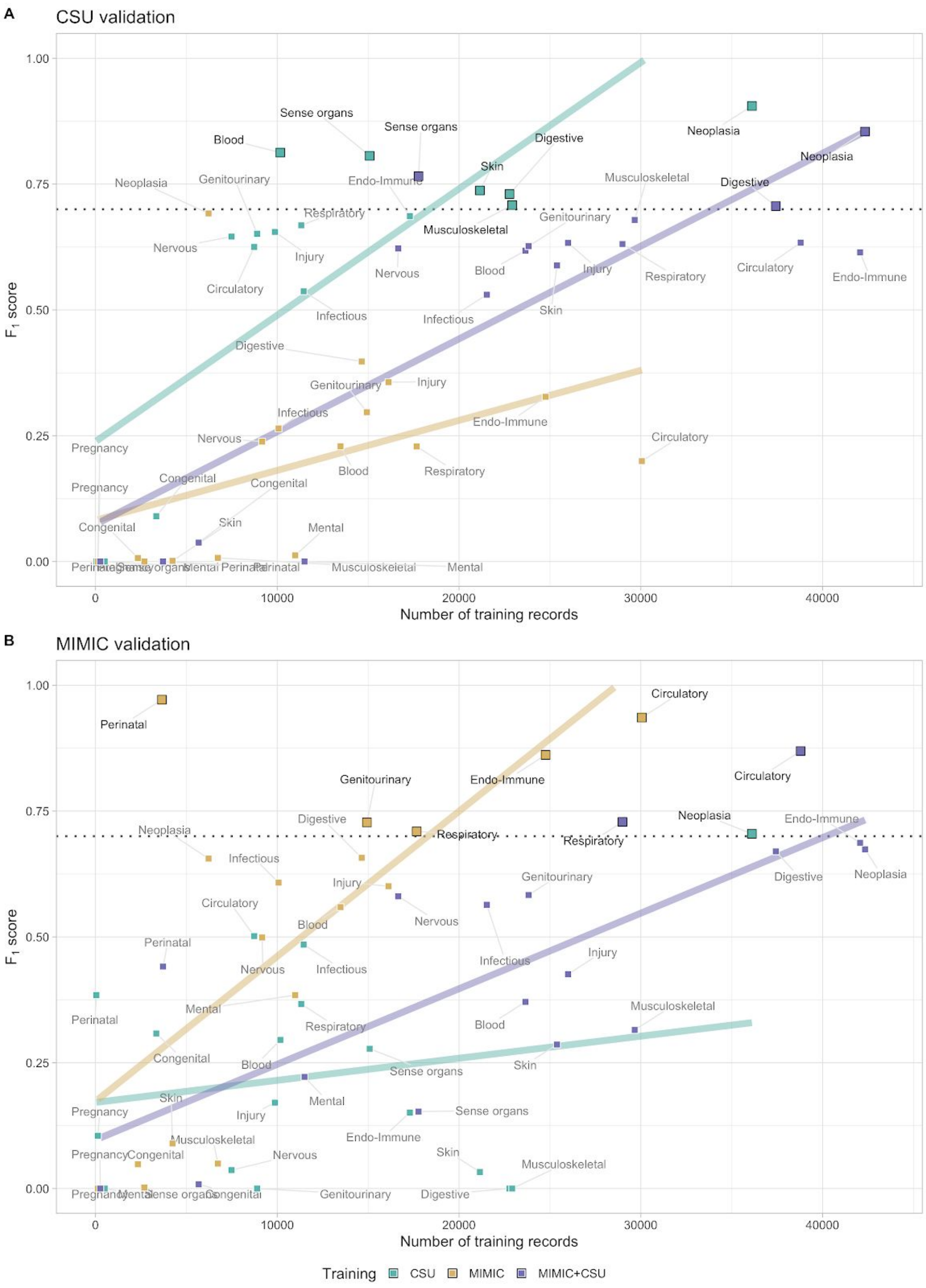


**Figure A**. $F_1$ scores by category using various training configurations. Training with CSU data (green), MIMIC data (yellow) or MIMIC+CSU (purple); validating on CSU data (Panel A) or MIMIC data (Panel B). The color of the category text is darkened (black) and the box made bigger if it surpasses the threshold of at least 0.70 $F_1$ score (dotted horizontal line).
